# Controllable Protein Design by Prefix-Tuning Protein Language Models

**DOI:** 10.1101/2023.12.03.569747

**Authors:** Jiawei Luo, Xianliang Liu, Jiahao Li, Yang Zhang, Qingcai Chen, Junjie Chen

**Author notes:** Corresponding author: Junjie Chen;.

## Abstract

The design of novel proteins with tailored functionalities, particularly in drug discovery and vaccine development, presents a transformative approach to addressing pressing biomedical challenges. Inspired by the remarkable success of pre-trained language models in natural language processing (NLP), protein language models (ProtLMs) have emerged as powerful tools in advancing protein science. While NLP leverages flexible text-based control tags to prompt language model generation, the restricted amino acid space (limited to 20 residues) imposes inherent constraints on achieving analogous controllability. In this study, we propose PrefixProt, a framework for controllable protein design that employs prefix tuning to learn virtual tokens as control tags. These virtual tokens are adaptively tailored to diverse protein properties through a data-driven manner and can be combinatorially integrated to enable multi-objective control over protein generation. The effectiveness of PrefixProt was validated through extensive experiments encompassing both protein structure design (e.g. alpha-helix or beta-sheet topologies) and protein function design (e.g. antimicrobial or anticancer peptide activities). Benchmark results demonstrate that prefix virtual tokens efficiently guide the pre-trained ProtLM by optimizing a smaller number of trainable parameters, out-performing other parameter-efficient fine-tuning methods and text-guided ProtLMs, particularly in scenarios with limited data availability. More importantly, the compositional flexibility of virtual tokens facilitates the generation of proteins with multiple target properties, substantially expanding the scope of design possibilities. By harmonizing controllability, efficiency and generalizability, PrefixProt establishes a robust framework for *de novo* protein design, with promising applications in drug discovery and biomedicine.

**Availability and implementation:** The models and code are available at: https://github.com/chen-bioinfo/PrefixProt

## 1 Introduction

Proteins perform a vast array of functions essential to biological processes. However, when repurposed for biomedical applications, such as drug discovery or vaccine design, they often suffer from a series of limitations, including structural instability, suboptimal geometry or size, low recombinant expression, and off-target interactions with other proteins [19]. These limitations stem from the biophysical constraints of naturally evolved sequences, underscoring the need for *de novo* protein design to systematically explore the vast amino acid sequence space and engineer bespoke functionalities. Advances in protein engineering over recent decades have yielded breakthroughs in enzyme catalysis [39], vaccine development [9], and therapeutic design [5], predominantly through experimental techniques like directed evolution [40] or computational approaches leveraging homology modeling and energy minimization [21, 22]. Nevertheless, such approaches are fundamentally constrained by evolutionary biases in natural protein repertoires or by their dependence on structural templates, limiting the exploration of novel sequence-structure landscapes beyond evolutionarily conserved motifs.

The availability of large-scale protein datasets has accelerated the development of artificial intelligence (AI)-driven protein design methods. Inspired by the remarkable success of transformer-based models in natural language processing (NLP), protein language models (ProtLMs) employ similar architectures to treat protein sequences as biological ‘languages’, enabling the decoding of sequence-function relationships inherent in natural proteins. To enhance controllability in protein generation, recent studies have adapted NLP paradigms like prompt learning [7, 23, 28], instruction learning [15] and parameter efficient fine-tuning (PEFT) [4, 16, 45] into ProtLMs. Prompt learning guides language models by prepending a prompt (a task-specific input template or cue) to generate contextually relevant outputs without modifying model’s parameters. In NLP, prompts are often natural language phrases (e.g., “Generate a poem”). However, protein vocabularies are restricted to 20 amino acids, preventing to directly construct text-like phrases [3] (**Figure 1 A**). To address this limitation, ProGen [35] encoded taxonomic and keyword tags, such as molecular function and cellular component, as special tokens to conditionally generate sequences. Nevertheless, such tags lack the flexibility to encode nuanced biochemical or structural constraints, providing limited control over protein generation. Instruction learning trains models to follow free-form natural language instructions (e.g., “Generate an anticancer peptide”) by fine-tuning on task-specific datasets containing input-output pairs. For instance, ProLLaMA [30], InstructProtein [47] and ProteinDT [27] integrated multi-modal text-protein pairs by employing instruction learning to align protein generation with free-text descriptors. However, challenges arise due to the disconnect between natural language instructions and the protein sequence space. Ambiguity in translating biochemical constraints (e.g., “improve binding affinity”) into actionable prompts reduces reliability. PEFT adapts a pre-trained model to a specific task by updating a subset of its parameters on a task-specific dataset [4]. As alternatives to full-parameter fine-tuning, PEFT mitigates computational cost and overfitting risks in low-data scenarios [31, 50]. For example, Low-Rank Adaptation (LoRA) [42, 48], which adds trainable low-rank matrices to ProtLM weights, has demonstrated resource-efficient advancements in protein function prediction tasks. Additionally, other PEFT methods, such as prefix-tuning [24], (IA)^3^ [26] and Vector-based Random Matrix Adaptation (VeRA) [20], were also widely applied in natural language generation tasks.

**Figure 1.**
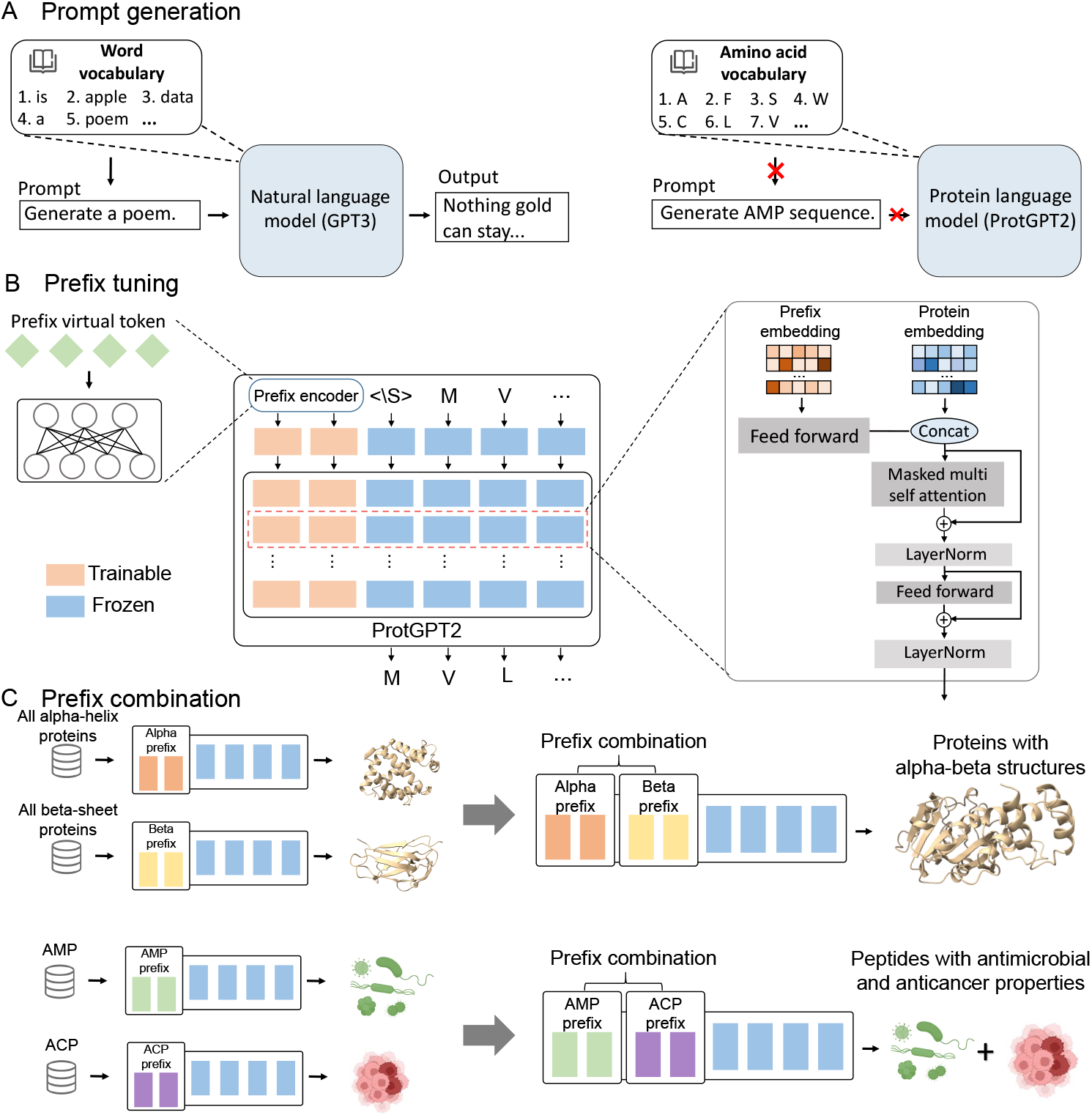
Illustration of the PrefixProt framework. (A) Prompt generation. Protein language models, constrained to 20 amino acids, lack flexible prompt engineering capabilities compared to natural language models. (B) PrefixProt framework. Prefix virtual tokens are prepended to protein sequences, steering amino acid generation downstream while preserving pre-trained parameters. (C) Multi-prefix control. Combinatorial integration of trained virtual tokens enables programmable generation of proteins with hybrid architectures (e.g. alpha-beta proteins) or dual-functional activities (antimicrobial and anticancer peptides).

Among PEFT paradigms, prefix-tuning is particularly different with other methods. It keeps the pre-trained model parameters frozen during optimization, while learning a compact task-specific prefix vector (referred to as prefix virtual tokens), composed of continuous and trainable embeddings. By treating this learned prefix as a virtual context that subsequent tokens can attend to, prefix-tuning effectively combines the adaptability of prompt-based learning strategies with the parameter efficiency inherent to PEFT frameworks. Specifically, prefix virtual tokens act as a task-specific embedding sequence that guides generation without altering the pre-trained model, enabling precise control over output properties while preserving computational scalability. As a result, prefix-tuning holds particular promise for controllable protein design.

In this study, we propose PrefixProt, a framework that leverages prefix-tuning to achieve controllable multi-property protein sequence design. PrefixProt learns virtual tokens as data-driven modular control tags, each encoding a specific property. These prefixes can be individually applied for single-property control or combinatorially integrated to generate proteins satisfying multiple-property constraints. We validate its effectiveness through extensive experiments encompassing both protein structural (e.g., alpha-helix and beta-sheet structures) and functional (e.g., antimicrobial and anticancer peptide functions) design tasks, demonstrating superior performance over PEFT methods and text-guided models in terms of success rate, structural validity and sequential realism of generated proteins, particularly in low-data scenarios. More importantly, PrefixProt enables compositional control, where combining virtual tokens generates proteins satisfying multiple constraints, thereby substantially expanding the scope of design possibilities, even to design the multi-property proteins that are not discovered in nature. By freezing the pre-trained ProtLM and optimizing only lightweight prefixes, PrefixProt combines the controllability of prompt engineering with the efficiency of PEFT, offering a scalable and resource-efficient paradigm for controllable protein design and advancing the frontier of AI-driven protein engineering.

## 2 RESULT

### 2.1 Overview of the proposed PrefixProt

PrefixProt is a parameter-efficient framework for controllable protein generation (**Figure 1**), designed to address the challenge of generating proteins with tailored multi-properties through prefix tuning on ProtLMs. While ProtLMs hold promising for *de novo* protein design, their utility in controllable protein generation remains constrained by the inherent limitations of their vocabulary (limited to 20 amino acid residues). This vocabulary precludes the use of natural language-like prompts for controllable generation.

The framework trains a lightweight prefix encoder to learn prefix virtual tokens. During training, the encoder generates these tokens through gradient optimization, while all pre-trained ProtLM parameters remain frozen. The prefix virtual tokens are prepended to protein sequences and dynamically interact with the frozen ProtLM’s transformer architecture. At each generation step, the concatenated token-sequence representations influence the autoregressive prediction of the next amino acid. This approach enables precise control over protein properties while maintaining computational efficiency, with trainable parameters accounting for merely 1% of the total model size. A key innovation of PrefixProt is its compositional control capability, where pre-trained prefix tokens act as a modular virtual language to define the target protein space. Each token semantically encodes biochemical constraints (e.g., structural topologies like alpha-helices or beta-sheets, or functional properties like antimicrobial/anticancer activity). By combinatorially integrating multiple tokens, the framework dynamically constructs hybrid specifications akin to composing linguistic phrases, enabling programmable generation of proteins with multi-constraint attributes.

We validate PrefixProt across two key axes: structural control (e.g., generating proteins with alpha-helical topology) and functional design (e.g., creating antimicrobial or anticancer peptides). Building on this foundation, we further demonstrate its capacity for compositional integration of multiple prefix virtual tokens to programmably generate proteins with hybrid properties. For instance, concatenating alpha-helix and beta-sheet (alpha-beta) tokens guides the design of chimeric architectures, while merging antimicrobial and anticancer (AM-AC) tokens generates multifunctional peptides. These tasks are infeasible for conventional methods, such as single-property fine-tuning (limited to isolated constraints) or text-guided ProtLMs (struggling with ambiguous natural language-to-sequence mappings), thereby highlighting the unique expressiveness of PrefixProt’s virtual token-based language.

### 2.2 Controllable generation of alpha-helical proteins

To evaluate the controllability of PrefixProt in generating structural and biophysical valid proteins, we benchmarked its performance in generating alpha-helical proteins against several baseline methods, including pre-trained ProtGPT2, fine-tuned Prot-GPT2, random generation based on the background distribution of amino acids, and three PEFT methods: LoRA [17], (IA)^3^ [26], VeRA [20], as well as two text-guided ProtLMs: ProLLaMA [30] and InstructProtein [47].Prefix virtual tokens of length 100, encoding alpha-helix structural constraints, were trained by PrefixProt using alpha-helix proteins from the Alpha class of the Structural Classification of Proteins (SCOP2) database, with trainable parameters accounting for only 1.18% of the total model.

#### Percentage of alpha-helical topologies

The proportion of amino acids in alpha-helical topologies in generated proteins was quantified using DSSP (**Figure 2A**). PrefixProt and LoRA achieved comparable performance with median alpha-helical contents of 60.8% and 61.0%, respectively, closely resembling those of natural alpha-helical proteins from the testing dataset, both significantly outperformed other methods. Among the PEFT methods, (IA)^3^ yielded slightly lower performance than prefix-tuning, while VeRA showed the weakest performance, but remained comparable to the fine-tuned model. The fine-tuned model significantly outperformed the pre-trained method. In contrast, ProLLaMA and InstructProtein performed poorly in generating proteins with alpha-helical structures, achieving results even worse than random generation, highlighting the inefficiency of text prompts in guiding structure-specific protein generation.

**Figure 2.**
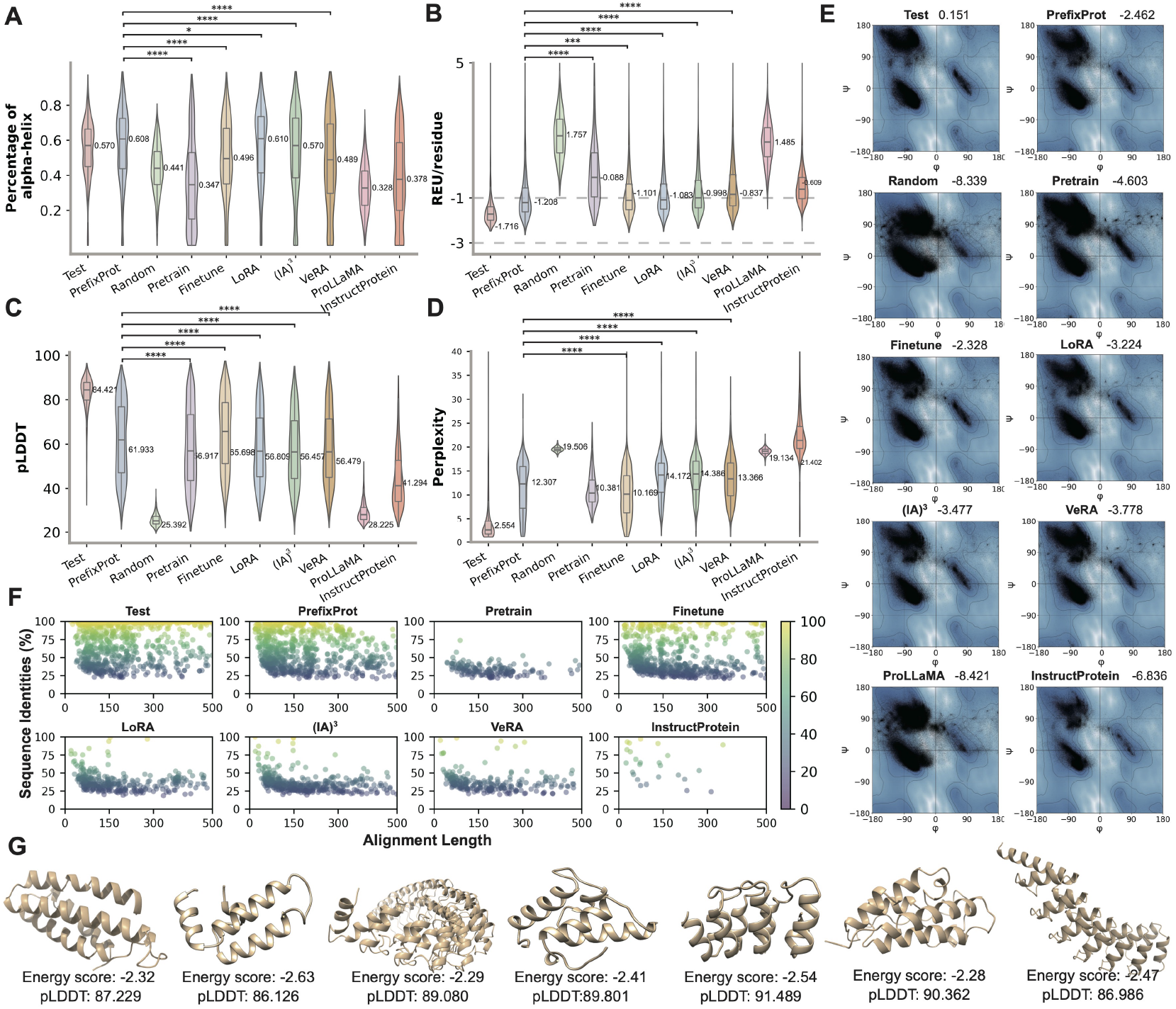
Comparison in generating proteins with alpha-helical structure. (A) Percentage of alpha-helix structure, (B) average Rosetta energy units per residue, (C) pLDDT of predicted protein structure, (D) perplexity of protein sequences, (E) Ramachandran plots of dihedral angular distribution quantified with Rama-Z score, (F) pairwise sequence identity against training dataset, and (G) seven alpha-helical proteins generated by PrefixProt. Significant levels: *∗* p*<*0.05, *∗∗* p*<*0.01, *∗ ∗ ∗* p*<*0.001, *∗ ∗ ∗∗* p*<*0.0001.

#### Structural stability and reasonability

To assess the structural validity of generated proteins, we calculated the Rosetta Energy Units (REU) scores and predicted Local Distance Difference Test (pLDDT) scores. The REU scores (**Figure 2B**) revealed that most of the generated proteins had energy scores between −1 and −3 REU/residue, except for those from the random generation. Proteins from the testing dataset showed a median REU score of −1.716 REU/residue, while those generated by PrefixProt, the fine-tuned model, and LoRA achieved median scores of −1.208, −1.101, and −1.083 REU/residue, respectively. Proteins generated by PrefixProt had lower Rosetta energy scores than those generated by the other methods, though they still higher than natural proteins in the testing dataset. Other PEFT methods performed comparably but with higher REU values. In contrast, the pre-trained ProtLM and text-guided ProtLMs performed poorly with positive REU values, particularly the performance of ProLLaMA was similar to random sequences, indicative of unstable or misfolded structures. The pLDDT results (**Figure 2C**) demonstrated that the fine-tuned model achieved the highest pLDDT scores with a median value of 65.70, followed closely by PrefixProt with a median pLDDT score of 61.93. Other PEFT methods (LoRA, (IA)^3^, and VeRA) achieved comparable scores around 56. In contrast, text-guided ProtLMs underperformed, where ProLLaMA exhibited the lowest performance similar with random sequences. To further demonstrate effective control over generation of alpha-helix proteins with stable conformations, the structures with energy score and pLDDT of seven all alpha-helix proteins generated by PrefixProt is visualized in **Figure 2G**. The conformational reasonability of generated proteins was further evaluated using dihedral angular (*ϕ, ψ*) distributions in Ramachandran plot [38] (**Figure 2E,** S1), where a Rama-Z score of *−*4 and lower indicates a serious problem with the structure. The proteins generated by PrefixProt and fine-tuned model achieved average Rama-Z scores with *−*2.462 and *−*2.328, indicating the protein backbone conformations generated by both of them were in favorable regions. In contrast, LoRA, (IA)^3^, and VeRA achieved average Rama-Z scores with *−*3.224, *−*3.477, and *−*3.778 showed significantly degraded performance. The other methods achieved Rama-Z scores lower than *−*4, indicating their protein backbone conformations have serious problems.

#### Sequence realism and diversity

The realism and diversity of generated protein sequences were assessed via perplexity and sequence identity. The perplexity results (**Figure 2D**) revealed that the fine-tuned model and PrefixProt achieved comparable performance with median perplexity scores of 10.17 and 12.31, outperforming PEFT methods. Same as previous metrics, text-guided ProtLMs (ProLLaMA and InstructProtein) exhibited the worst performance similar with random sequences. Sequence identity was calculated by aligning generated proteins against the training dataset, where higher identity scores indicate the generate proteins are similar with proteins in training dataset (**Figure 2F**). The proteins produced by PrefixProt exhibited a widely range of sequence identity scores across various sequence length (less than 500 amino acids), similar to the testing dataset. These results indicate that PrefixProt effectively captured the biological patterns in natural proteins, while maintaining the capacity to explore the novel protein sequences with alpha-helical structures. In contrast, the proteins generated by fine-tuned model and PEFT methods (LoRA, (IA)^3^, VeRA) predominantly exhibited sequence identity scores falling within the range of 20% to 50%, indicating that the generated protein sequence deviates from natural patterns. Text-guided methods exhibited extreme divergence, further highlighting their inability to capture natural protein patterns. The results for diversity and novelty values are presented in Table S1. PrefixProt achieved a moderate novelty score of 149.39, surpassing text-guided methods, while PEFT methods LoRA, (IA)^3^, and VeRA obtained 170.83, 156.98, and 168.35, respectively. Additionally, PrefixProt demonstrated competitive diversity (141.26), outperforming text-guided approaches (ProLLaMA: 132.82, InstructProtein: 50.87) as well as (IA)^3^ (136.68) and VeRA (124.43). These results indicated that PrefixProt achieved a good balance between diversity and novelty without sacrificing structural specificity.

Overall, PrefixProt, with a limited number of trainable parameters, outperformed the compared methods in generating proteins with tailored alpha-helical structures, stable energy, reasonable conformation and diverse sequence identity. These results demonstrate the capability of PrefixProt to effectively prompt large ProtLM toward generating proteins with desired structural properties.

### 2.3 Controllable generation of functional peptides

To evaluate the capability of PrefixProt in generating proteins with tailored functions, we benchmarked its performance in generating peptides with antimicrobial or anticancer activities against PEFT methods (LoRA, (IA)^3^ and VeRA) and text-guided ProtLMs (ProLLaMA and InstructProtein). Besides, for AMP generation, PrefixProt was also compared with three specialized AMP design methods: PepLSTM [33], PepCVAE [10], and HydrAMP [43]. Two task-specific virtual tokens of length 20 encoding AMP or ACP properties were learned by PrefixProt on DBAASP and AntiCP2.0 datasets, with trainable parameters accounting for only 0.24% of the ProtLM model.

#### Percentage of functional peptides

Functional probabilities of generated AMPs and ACPs were predicted using CAMP [46] and AntiCP2 [1], respectively (**Figure 3A,D**). PrefixProt achieved the highest performance with median probability scores of 0.90 (AMP) and 0.89 (ACP), while LoRA achieved comparable performance with 0.89 (AMP) and 0.88 (ACP), both significantly outperforming other PEFT methods. While text-guided models (ProLLaMA: 0.79 AMP, 0.045 ACP; InstructProtein: 0.35 AMP, 0.09 ACP) underperformed even compared to random sequences. Notably, VeRA and text-guided ProtLMs (ProLLaMA and InstructProtein) showed extremely low on ACP generation, which may be due to limited data availability of ACPs. PrefixProt also significantly outperformed existing specialized AMP design models (PepLSTM: 0.80, PepCVAE: 0.74, and HydrAMP: 0.67). These results suggest that PrefixProt has strong potential for generation of functional peptides, compared with the conventionally specialized generative methods.

**Figure 3.**
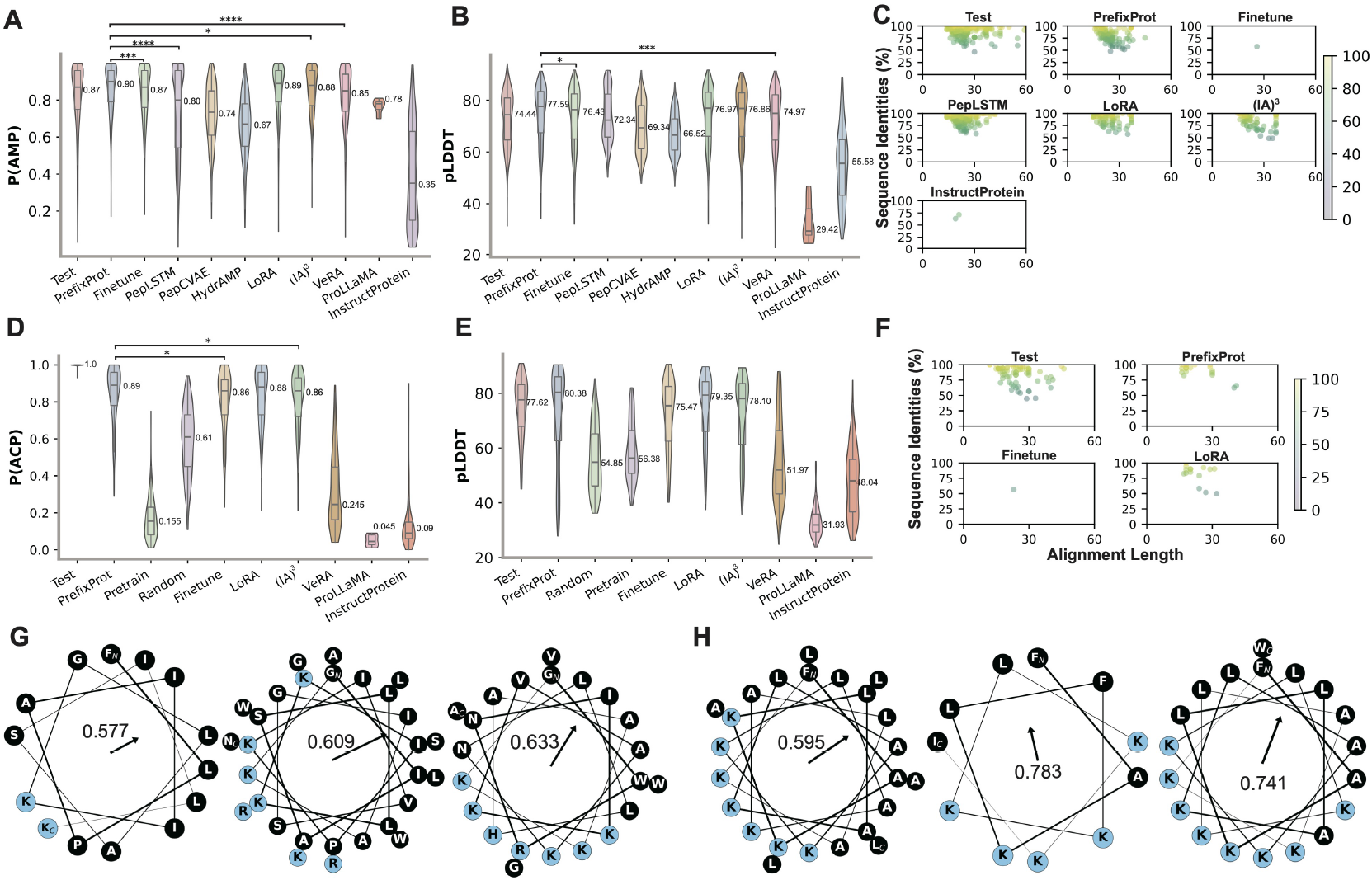
Comparison in generating proteins with antimicrobial or anticancer functions. The probability distribution comparison of being AMPs (A) and ACPs (D), pLDDT of predicted peptide structure comparison of AMPs (B) and ACPs (D), pairwise sequence identity comparison of AMPs (C) and ACPs (F), and the helical wheel plots of 3 top-ranking AMPs (G) and ACPs (H) generated by PrefixProt. The amino acids labeled with ‘N’ and ‘C’ subscript represent the N-terminus and C-terminus of the peptide sequences, respectively. The black and blue circles represent hydrophobic and polar residues, respectively. The numbers and corresponding arrows depict Eisenberg’s hydrophobic moment for the idealized *α*-helical conformation. Significant levels: *∗* p*<*0.05, *∗∗* p*<*0.01, *∗ ∗ ∗* p*<*0.001, *∗ ∗ ∗∗* p*<*0.0001.

#### Structural validity and amphipathicity

The structural validity of generated AMPs and ACPs was measured with pLDDT scores (**Figure 3B,E**). PrefixProt achieved the highest performance with median pLDDT scores of 77.59 (AMP) and 80.38 (ACP), while PEFT methods and the fine-tuned model achieved comparable performance. In contrast, text-guided ProtLMs, especially ProLLaMA, exhibited the lowest structural confidence, failing to form reasonable conformations. The amphipathicity of AMPs and ACPs generated by PrefixProt was assessed using the helical wheel plots of the top 3 AMPs ranked by predicted antimicrobial or anticancer probability respectively (**Figure 3G,H**). Amphipathicity is a significant property influencing the antimicrobial activity [13], as shown by spatial segregation between hydrophobic and polar residues. This segregation can be quantified using the hydrophobic moment (HM) [11], where the HM vector points toward the hydrophobic face of the helix. The helical wheel plots clearly illustrated that the amphipathic distributions of AMPs and ACPs exhibit favorable amphipathicity characteristics.

#### Sequence diversity and physicochemical fidelity

The sequence identity for generated AMPs and ACPs was conducted against the corresponding training dataset (**Figure 3C, F**). AMPs and ACPs produced by PrefixProt, PepLSTM exhibited varied sequence identities ranging from 50 to 100 to those in training datasets, similar as the natural proteins in test, demonstrating that they have effectively captured the biological patterns in natural AMPs or ACPs, while maintaining the capacity to explore the novel sequences space. In contrast, LoRA and (IA)^3^ generate fewer matched sequences compared to PrefixProt, with only one sequence produced by the fine-tuned ProtGPT2 matching the training set. Regarding to the three specialized AMP design methods (HydrAMP, PepLSTM, and PepCVAE), only PepLSTM generated sequences with comparable patterns. The sequences generated by PepLSTM predominantly exhibited over 70% similarity to known peptides but were notably shorter in length, suggesting its limitation in generalization ability. In contrast, HydrAMP and PepCVAE failed to identify any discernible patterns, which might stem from significant discrepancies between their training protocols and the alignment datasets used for evaluation. The generated peptides were then compared in terms of 7 physicochemical properties, including amino acid distribution, length, charge, charge density, hydrophobicity, hydrophobic moment, and isoelectric point (**Supplementary Figure S2, S3**). Except for AMPs generated by pre-trained ProtGPT2 and text-guided ProtLMs, all other methods are able to produce AMPs and ACPs with amino acid distribution close to natural peptides in the test dataset. Note that the random generation is based on the background distribution of amino acids in the testing dataset. Thus it has the same length distribution with training data. The other results of 6 physicochemical property distributions show that the AMPs and ACPs generated by PrefixProt, fine-tuned ProtGPT2 and PEFT methods(LoRA, (IA)^3^, VeRA) are notable resemblance with those of natural AMPs and ACPs in the testing dataset respectively.

Overall, PrefixProt outperformed the compared methods in generating proteins with tailored antimicrobial function and anticancer function. PrefixProt was even better than the specialized AMP design methods. These results demonstrate PrefixProt has the capability to effectively prompt large ProtLM toward generating proteins with desired functional properties.

### 2.4 Combing virtual tokens for multi-property generation

PrefixProt enables compositional control over protein generation by integrating learned virtual tokens to impose multi-objective constraints. This capability was evaluated through two design tasks: 1) generation of hybrid structural proteins containing both alpha-beta conformations and 2) design of dual-functional peptides with concurrent AM-AC activities. Two combination strategies were explored: concatenation (PrefixProt-concat), which preserves individual token embeddings) and averaging (PrefixProt-avg), which blends embeddings into a unified representation. And they were compared with baseline methods, including PEFT (prefix tuning, LoRA, (IA)^3^, VeRA) and Finetune (with full parameters). The compared baseline methods were trained on natural multi-property proteins.

### 2.4.1 Hybrid structural protein generation

To generated hybrid structural proteins (alpha-beta proteins), distinct virtual tokens encoding alpha-helix and beta-sheet conformations were pretrained, each with 20 tokens due to the PortLM’s input length constraints. The compared baseline methods, including PEFT and Finetune, were trained on natural alpha-beta proteins. Since the PEFT methods are parameter-efficient methods that require low-data, we set a small dataset with randomly selected 500 alpha-helix proteins, 500 beta-sheet proteins, and 500 alpha-beta proteins. Generated sequences were structurally validated using ESM-Fold predictions, with quality assessed via pLDDT and Rama z-scores.

As illustrated in **Figure 4A**, PrefixProt-concat and PrefixProt-avg achieved median pLDDT scores of 55.20 and 55.21, respectively, outperforming PrefixProt, LoRA and (IA)^3^, while marginally underperforming VeRA and Finetune methods. The energy score (**Figure S4A**), perplexity (**Figure S4B**) and sequence identity (**Figure S4D**) also demonstrated that, under the prompt of combined alpha-beta tokens, ProtLM can still hold on the ability to generate high-quality proteins. The further analysis of Rama z-scores in the high-quality structures (pLDDT*>*70) revealed that all methods achieved Rama z-scores in range of −1.0 ~ −1.6, indicating the generated protein backbone conformations were in favorable regions (**Figure 4D, S4C)**.

**Figure 4.**
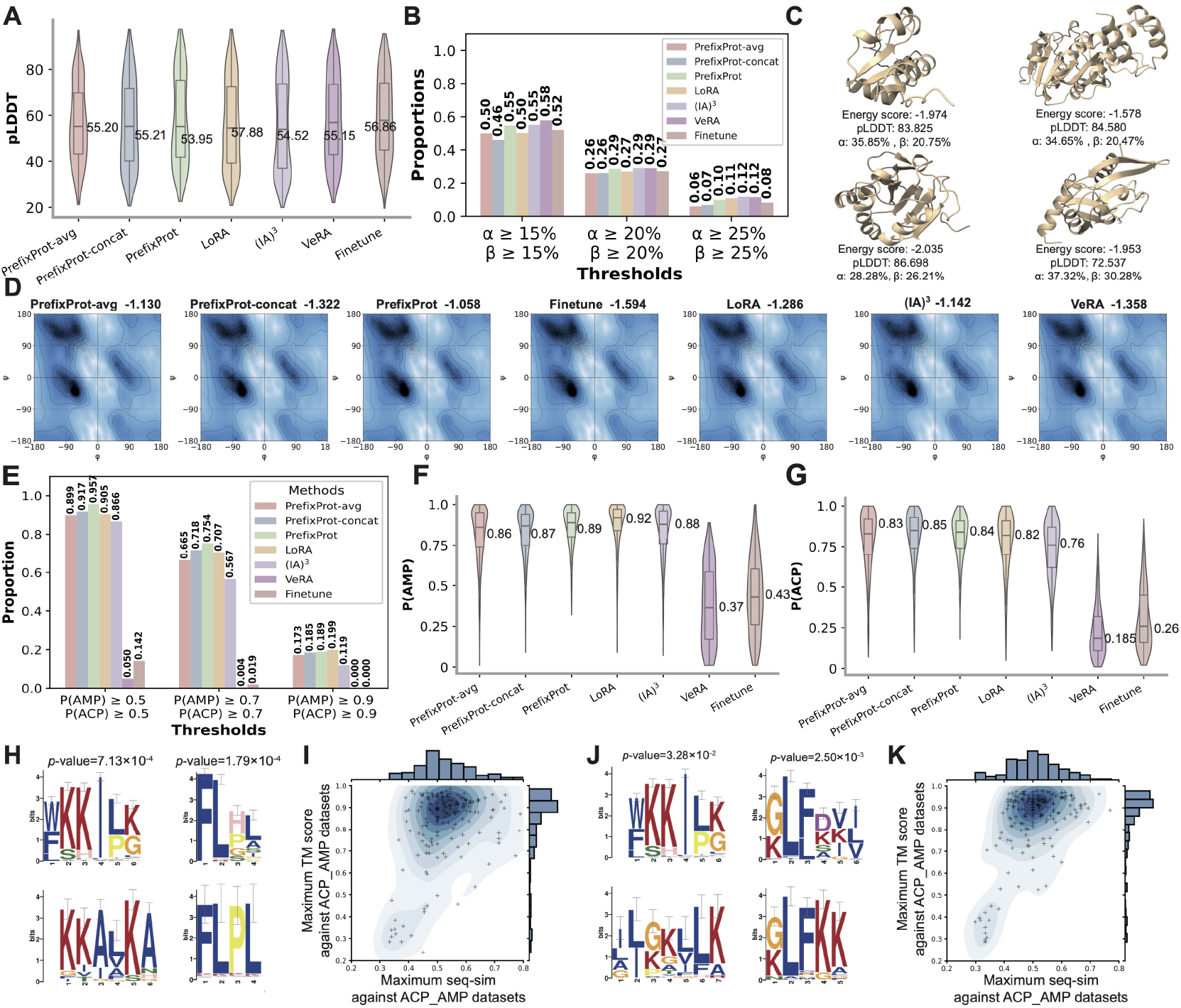
Comparison of multi-properties controllable generation. (A) pLDDT distribution of predicted protein structure. (B)Proportion of high-confidence generated proteins (pLDDT*>*70) with alpha-helix and beta-sheet content exceeding defined thresholds of 0.15, 0.20, and 0.25. (C) Four alpha-beta protein structures generated using averaging (top) and concatenation (bottom) prefix combination approaches. (E) Proportion of generated AM-AC peptides with activities exceeding probability thresholds of 0.5, 0.7, and 0.9. (F, G) Probability distribution of being AMPs (F) and ACPs (G). (H, J) Motif comparison for natural AM-AC peptides (top) and generated AM-AC peptides (bottom). (I, K) are the distribution of sequence similarity (seq-sim) and structural similarity (TM-score) between the natural and generated AM-AC peptides using averaging (I) and concatenation (K) prefix combination approaches.

To assess structural composition, alpha-helix and beta-sheet content were quantified at thresholds of 15%, 20% and 25% (**Figure 4B**). More than 50% of high-quality protein structures generated by PrefixProt-concat and PrefixProt-avg exceeded 15% content for both alpha-helix and beta-sheet conformations. While their performance was slightly worse than other compared methods at the thresholds of 15%, PrefixProt-concat and PrefixProt-avg had comparable performance with other compared methods at thresholds *≥*20%. Notably, PrefixProt-concat marginally outperformed PrefixProt-avg at lower thresholds (15%), but difference diminished at higher thresholds (*≥*20%). Four representative generated alpha-beta proteins by PrefixProt-concat and PrefixProt-avg exhibited high and balanced alpha-beta content with favorable energy and pLDDT scores (**Figure 4C**). The results on large dataset were shown in **Supplementary Figure S4E,F**

Critically, PrefixProt that combined pretrained tokens without additional training on alpha-beta proteins has the capability to generate proteins with hybrid structural features even under data-constrained conditions. The two combination strategies have no significant difference in generation of hybrid structural proteins with alpha-helix and beta-sheet conformations. This further underscores its effectiveness and flexibility in producing structurally valid and compositionally complex proteins.

#### 2.4.2 Dual-function peptides design

To isolate the impact of token combination, antimicrobial tokens and anticancer tokens were pretrained on only AMPs and only ACPs datasets, respectively, excluding 353 natural AM-AC peptides. The generated AM-AC peptides were functionally validated using CAMP [46] and AntiCP 2.0 [1] predictors. PrefixProt-concat and PrefixProt-avg achieved high dual-function proportions above the probability thresholds of of 0.5 and 0.7 (**Figure 4E**), surpassing LoRA and (IA)^3^ methods, though slightly worse than PrefixProt (trained on natural AM-AC peptides). Finetune and VeRA were even failed to generate the AM-AC peptides. Regarding to the separately antimicrobial or anticancer activity distributions of the generated peptides (**Figure 4F, G**), both PrefixProt-concat and PrefixProt-avg also achieved superior performance, even better than PrefixProt in distribution of probability of ACP. Besides, PrefixProt-concat is slight better than PrefixProt-avg. These findings demonstrate that the tokens combination can effectively generate multi-functional proteins, and the concatenation manner has higher controllability than the average manner.

We then evaluated the captured patterns and sequence novelty by comparing the motifs and sequence-structure similarity against natural AM-AC peptides. The motifs were analyzed via STREME (**Figure 4H, J**) with the parameters in **Supplementary Table S2**. Both PrefixProt-concat and PrefixProt-avg captured the motif ‘WKKILK’, but they also have different sequence patterns, suggesting that the two combination strategies have distinct effects on generation progress. Moreover, the distribution of sequence-structure similarity revealed that the generated high probability AM-AC peptides (*≥* 0.9) exhibited low sequence similarity (*~*50%) while maintaining high structural similarity (*~*0.9 TM-scores) to natural AM-AC peptides (**Figure 4I, K**), demonstrating that the generated AM-AC peptides are novel in their amino acid sequences while remaining effective in functional activity. Pairwise sequence identity (**Figure S5A**) results also showed that the generated sequences exhibited high diversity, indicating that the model was not simply memorizing but creating novel variants. The structure alignments between generated AM-AC peptides and natural AM-ACPs further validated functional conservation (**Figure S5B**).

While the concatenation strategy achieved higher performance in hybrid alpha-beta structure proteins and dual AM-AC function peptides design, it need more parameter costs due to extended prefix lengths, wheres the averaging combination does not alter the prefix length. This highlights a trade-off between controllability and parameter efficiency. By combining virtual tokens, PrefixProt enables multi-property protein generation without additional training, demonstrating flexibility in designing structurally and functionally complex biomolecules. This framework establishes a foundation for extending token-based control to higher-order protein engineering tasks.

### 2.5 Impact of training data scarcity and prefix length

To evaluate the robustness of PrefixProt under data-limited scenarios, we systematically evaluated its performance randomly by subsampling the full AMP dataset to create 6 small datasets containing 50, 100, 200, 500, 1000, and 2000 AMPs (**Figure 5B**). We employed the pre-trained ProtGPT2 as a compared baseline. The peptides generated by ProtGPT2 exhibited no antimicrobial activity, whereas PrefixProt significantly enhanced the capability of generating AMPs from a median *P*_*AMP*_ of 0.09 to 0.68 on only 50 AMPs, indicating its efficiency on extreme low-data constraints. As the training data size increasing, the success rate rose to a median *P*_*AMP*_ of 0.90 when using all 3604 AMPs.

**Figure 5.**
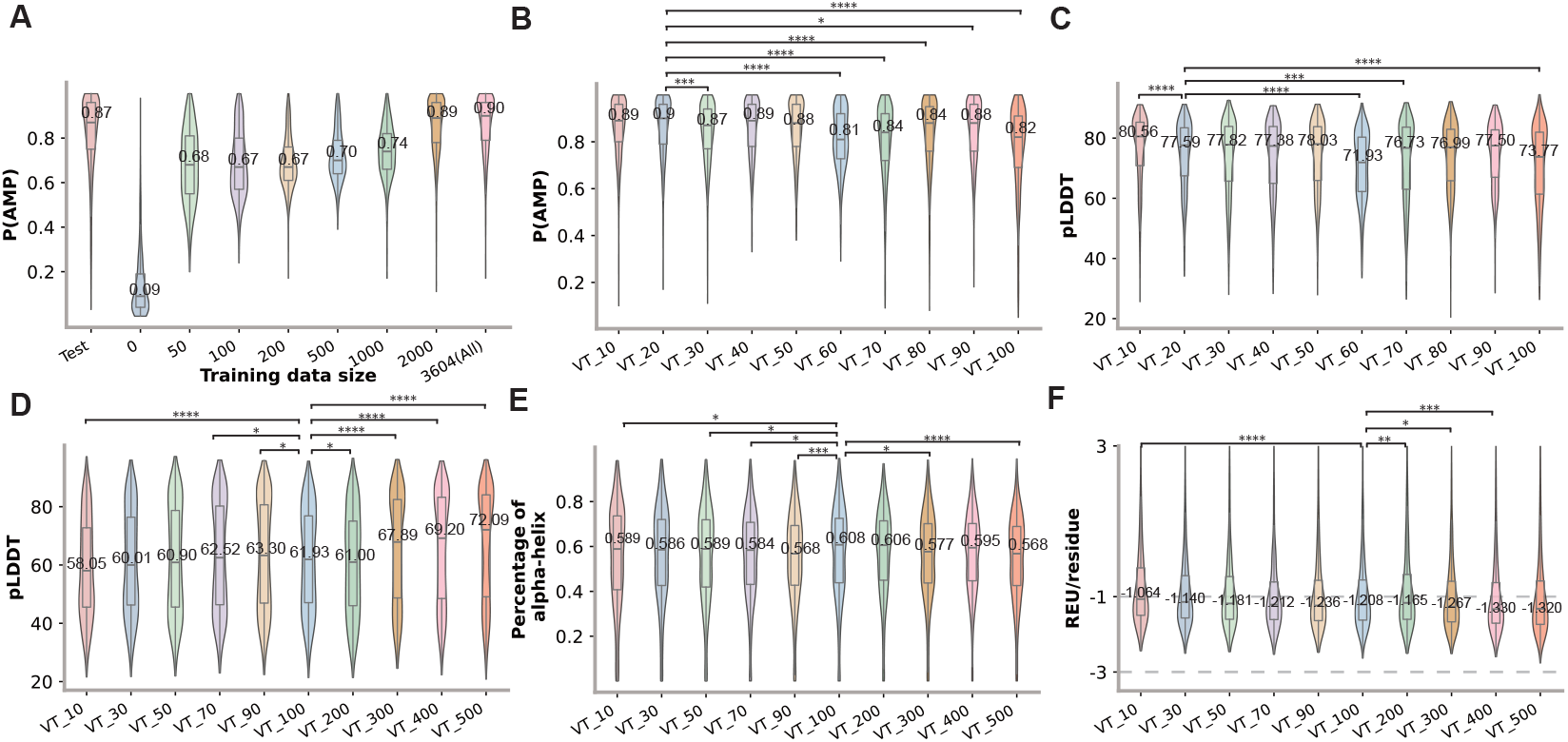
Comparison of antimicrobial activity in low-data and varying prefix length settings. (A) Probability distribution of AMPs generated by PrefixProt with token length of 20 across various low-data settings. (B and C) Probability distributions for AMPs and pLDDT scores generated by PrefixProt with different prefix lengths in the full AMP dataset. (D, E, and F) Percentage of alpha-helix structure, average Rosetta energy units per residue, and pLDDT scores generated by PrefixProt with different prefix lengths in the full alpha-helical protein datasets. Significant levels: *∗* p*<*0.05, *∗∗* p*<*0.01, *∗ ∗ ∗* p*<*0.001, *∗ ∗ ∗∗* p*<*0.0001.

We further investigated the impact of prefix token length on model performance using the AMP dataset (3604 sequences) and alpha-helical dataset (13,750 sequences) (**Figure 5C-F**). The AMP token length is progressively increasing from 10 to 100 due to small size of AMP dataset, and the alpha-helical token length is progressively increasing from 10 to 500. As the AMP token length increasing, the generated AMP activity has not significantly changing, even became slightly worse with the longer tokens due to more parameters are under trained. In contrast, the longer alpha-helical tokens significantly enhance the quality of generated alpha-helical proteins in terms of energy score (REU), pLDDT and perplexity. These results demonstrated that longer prefixes enhance expressivity but demand larger training set. These results highlighted that longer prefixes enhance model expressivity but require larger training datasets. Thus, for limited-data scenarios, short prefixes (e.g., 10-20 tokens) are sufficient, whereas longer prefixes should be explored for large-scale datasets.

## 3 Conclusion

This study introduced PrefixProt, a novel framework for controllable protein design that leverages prefix tuning to guide pre-trained ProtLMs through learned virtual tokens as property control tags encoding specific functional or structural properties. By freezing pre-trained parameters while exclusively training lightweight prefix embeddings, PrefixProt achieved superior performance in generating proteins with tailored alpha-helix conformations, antimicrobial/anticancer activity. Crucially, the method enables combinatorial integration of multiple virtual tokens, facilitating multi-property control (e.g., alpha-beta hybrid-structural topologies, dual-functional AM-AC peptides) without reliance on multi-property training datasets. This approach circumvents the need for extensive retraining, significantly enhancing computational efficiency and accessibility. PrefixProt addresses critical challenges in AI-driven protein engineering, offering transformative potential for biomedical applications, such as: 1) high-throughput design of multifunctional biomolecules (e.g. antibodies or dual-targeting cytokines with combined antiviral and anti-inflammatory properties, and 2) personalized therapeutic design through combinatorial token tuning aligned with patient-specific biomarkers. By enabling resource-efficient, multi-objective design, PrefixProt democratizes access to advanced protein engineering, particularly for academic and small-scale facilities with limited computational or experimental resources. Although its robustness, several limitations require further investigation. First, the efficacy of prefix combinations depends on token length and dataset scale, necessitating task-specific optimization; Second, the learned prefix embeddings limited interpretability and may potentially encode unintended noise-derived constraints that could bias generation outcomes.

Future work will focus on the library of prefix tokens to encode finer structural and functional properties, as well as developing systematic strategies for their combinatorial integration. Besides, it also could be interesting on integrating multi-modal inputs (e.g. structural embedding from AlphaFold, functional descriptors from text tokens) to enhance cross-domain controllability across sequence, structure and function. Additionally, adapting prefix tuning to chemical molecular design with SMILE representation could further broaden the framework’ applicability.

## 4 Methods

### 4.1 Datasets

To systematically evaluate the efficacy of prefix tuning in guiding ProtLMs for structure and function aware protein generation, we constructed alpha-helix and beta-sheet structural datasets and antimicrobial and anticancer functional datasets. For all datasets, 80% of the data was randomly sampled as training data, and the remaining 20% was the test data.

#### Alpha-helix and beta-sheet structural datasets

The alpha-helix proteins were derived from the Alpha class of Structural Classification of Proteins extended (SCOPe) database [14], which integrates manual annotations and computational methods to provide a comprehensive description of the structural and evolutionary connections among known proteins. After eliminating duplication, non-protein sequences, and sequences with non-standard amino acids, the final dataset contains 9,168 high confidence alpha-helix rich protein sequences, 14,531 beta-sheet protein and 14,223 alpha-beta proteins.

#### Antimicrobial and anticancer functional datasets

The antimicrobial peptides, a class of small proteins with short amino acid sequences that possess broad-spectrum antimicrobial activity for the treatment of drug-resistant bacterial infections, were sourced from the Database of Antimicrobial Activity and Structure of Peptides (DBAASP) [6, 36]. Sequences were classified as active based on experimentally validated inhibitory concentrations satisfying at least one IC50 *≤*10 *µ*M for *P. aeruginosa, ≤*10,000 nM for *A. baumanii*, or 32 *µ*gmL^*−*1^ for *S. aureus*. After quality control, 4,505 active AMPs were retained. The anticancer peptides utilized in this study were sourced from AntiCP2.0 [1], a curated database containing 861 experimentally validated anticancer peptides. This dataset was derived through rigorous processing of the original CancerPPD database, which involved the removal of non-natural sequences, short peptides (*<*5 amino acids), long chains (*>*50 amino acids), and redundant identical sequences. To benchmark the dual-functional generation, we constructed a composite dataset by intersecting sequences from AMP and ACP datasets and supplementing with data from MLBP [44], resulting in 353 dual-functional (AM-AC) peptides. In the prefix concatenation experiments, during the pretraining of ACP tokens and AMP tokens, known AM-AC peptides were excluded from the datasets. The AMP tokens were re-trained on 4,253 AMP-specific sequences, while the ACP tokens utilized 609 ACP-specific sequences, thereby minimizing functional overlap during model training.

### 4.2 Autoregressive Language Models

Autoregressive language models generate sequences by predicting each token conditioned on its previous context. For a protein sequence *a* = (*a*_1_, …, *a*_*L*_) of length *L*. The chain rule of probability can be formulated as 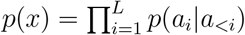 [2]. Autoregressive ProtLMs learn to capture evolutionary patterns and structural constrains in protein sequences through this sequential factorization. In practically, a ProtLM with parameters *θ* are optimized by minimizing the negative log-likelihood loss value across a given protein dataset *D* = *{a*^1^, …, *a*^|*D*|^*}*. The loss function can be formulated as

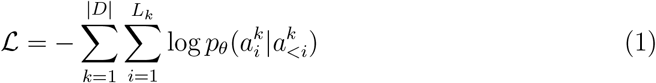

where *L*_*k*_ denotes the length of the *k*-th sequence.

Our framework employed ProtGPT2 [12] as the base model for protein generation. ProtGPT2 adopts GPT2-large architecture [37], comprising a 36 transformer decoder layers with 1280-dimensional embeddings and 12 attention heads in each layer. The self-attention mechanism operates through three learnable projections of input embeddings 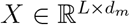 :

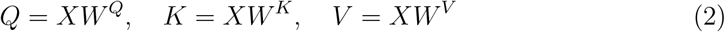

where 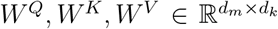 are learnable parameters. The scaled dot-product attention with causal masking ensures autoregressive constraints:

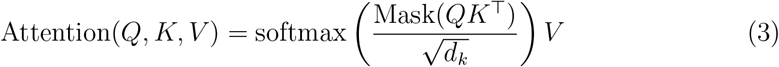

To capture diverse representations, multi-head attention executes *h* parallel attention heads:

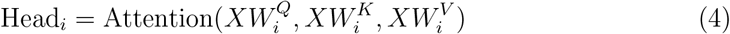

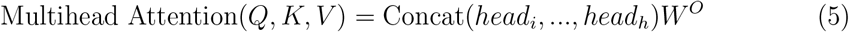

where the 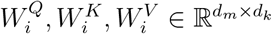 are the head-specific projections and 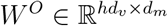 combines outputs from *h* heads. This architecture enables hierarchical learning of protein sequences features while maintaining generation coherence through autogressive constraints.

### 4.3 Parameter efficient fine-tuning

Parameter Efficient Fine-Tuning (PEFT) is a set of techniques designed to adapt large pre-trained machine learning models to specific tasks while minimizing the number of parameters that need to be updated. This approach addresses the computational and memory challenges associated with fine-tuning massive models, which often have billions of parameters. In this study, we employed prefix tuning to construct the proposed framework, and compared with LoRA, (IA)^3^, VeRA.

Prefix tuning adds trainable vectors to the input sequence, influencing model behavior via attention mechanisms. In this study, prefix tuning was employed to learn a virtual token as continuous prompts to steer ProtLMs toward generating proteins with desired functional and structural properties. Unlike discrete amino acid tokens, the prefix tokens are task-specific embedding optimized through gradient descent. Formally, given a protein sequence *a* and prefix *p* with length |*p*|, the prefix *p* is concatenated on the left of the protein sequence as the first context, thereby influencing the generation of amino acids on its right. The augmented input becomes *a*^*′*^ = [*p*; *a*], where *p*_*i*_ denotes the *i*-th virtual token. The prefix embeddings are generated by the prefix encoder *P*_*β*_ *∈* ℝ^|*p*|*×d*^, enabling conditional generation through context modulation. The hidden state *h*_*i*_ at position *i* can be formulated as:

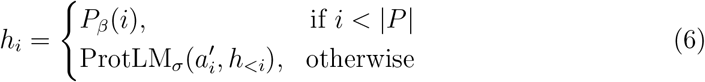

where *σ* and *β* represent ProtLM and prefix encoder parameters, respectively. Furthermore, to stabilize optimization, each *P*_*β*_[*i*, :] is reparameterized via *MLP*_*β*_(*Pβ′* [*i*, :]), where *Pβ′* is a trainable matrix. During training, the ProtLM parameters *σ* remains frozen, while only the parameters *β* in the prefix encoder are trainable, reducing memory overhead.

Low-Rank Adaptation (LoRA) [17] adapts a pre-trained model to a new task or domain by introducing low-rank matrices that modify the model’s weights, rather than updating all the parameters of the model. For a pre-trained weight matrix of ProtLM *W*_0_ *∈* ℝ^*d×k*^, we constrain its update by representing the latter with a low-rank decomposition:

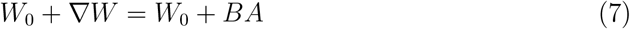

where *B ∈* ℝ^*d×r*^, *A ∈* ℝ^*r×k*^, and the rank *r ≪* min(*d, k*). Here, *A* and *B* are trainable low-rank matrices initialized via zero-centered Gaussian distributions, while *W*_0_ remains fixed. This constrains the update rank to *r*, reducing trainable parameters by *O*(*dr*+*rk*) compared to full fine-tuning. For autoregressive protein generation, LoRA applies this decomposition to attention and feed-forward layers while preserving the original next-token prediction objective.

Infused Adapter by Inhibiting and Amplifying Inner Activations (IA)^3^ [26] injects learnable scaling vectors into specific layers of a pre-trained model. These vectors rescale intermediate activations (e.g., attention keys/values or feed-forward outputs) to steer the model’s behavior for a target task. Formally, for an activation tensor *X ∈* ℝ^*T ×d*^, three task-specific learnable vectors *s*_*q*_, *s*_*k*_, *s*_*v*_ *∈* ℝ^*d*^ modulate queries, keys and values in self-attention:

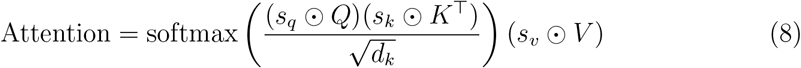

where *⊙* denotes element-wise multiplication. Similarly, the vector *s*_*ff*_ *∈* ℝ^*ff*^ scales activations in feed-forward networks (*s*_*ff*_ *⊙ γ*(*W*_1_*x*))*W*_2_, where *γ* denotes nonlinearity in the feed-forward network. Each Transformer layer incorporates independent scaling vectors, introducing *L*(*d*_*q*_ + *d*_*k*_ + *d*_*v*_ + *d*_*ff*_) parameters for *L* layers. All these vectors are initialized to ones to ensure stable training and preserve pretrained knowledge, with only 0.01% of the total parameters updated. This makes (IA)^3^ particularly suited for low-data structural/functional protein design.

Vector-based Random Matrix Adaptation (VeRA) [20] decomposes low-rank adaptation into fixed random matrices and trainable scaling vectors. For a weight matrix *W∈* ℝ^*d×k*^, the adaptation Δ*W* is computed as Δ*W* = *A·*diag(*b*) *·B*, where *A ∈* ℝ^*d×r*^ and *B∈* ℝ^*r×k*^ are frozen Gaussian random matrices shared across tasks, while *b ∈*ℝ^*r*^ is learnable task-specific. This reduces parameter count to *O*(*r*) per matrix, enabling multi-task adaption with minimal storage. VeRA’s fixed bases prevent overfitting while allowing functional specialization through learned scaling coefficients.

### 4.4 Text-guided protein language model

Text-guided protein language model integrates natural language instructions with protein sequence generation or optimization, enabling researchers to design or modify proteins using descriptive text prompts. This approach leverages advances in large language models (LLMs) and generative AI to bridge natural language understanding with biological sequence design. In this study, we introduced two text-guided ProtLMs as the baseline methods.

ProLLaMA builds on LLaMA2, enhanced through a two-stage training pipeline to harmonize protein sequence modeling with natural language comprehension. First, it continues pre-training from LLaMA2 on a large corpus of protein sequences, thereby preserving the model’s natural language capabilities while learning protein language. Second, instruction tuning was performed on a newly constructed dataset containing over 13 million task-oriented samples, such as controllable protein generation and protein superfamily prediction. For baseline generation, task-specific prompts were formulated using superfamily annotations: (1) For alpha-helix protein generation: Superfamily =*<*Single alpha-helix domain superfamily*>*; (2) For antimicrobial or anticancer peptide generation: Superfamily=*<*Short peptides with antimicrobial activity*>* and Superfamily=*<*Short peptides with anticancer activity*>*.

InstructProtein introduces a cross-modal strategy to align protein language with human language. It was first pretrained on a hybrid corpus, establishing a dual-language foundation. Next, a protein knowledge graph was constructed from heterogeneous sources, capturing their properties, families, and functions. Then, these graph triples are transformed into instructional templates, each containing a task description (in human language), an input (protein text or sequence), and the corresponding output. At last, InstructProtein used this supervised instruction set to align protein and human language representations. In this study, the text-based instructions for alpha-helix protein generation instruction is: I would like to generate a protein that has all-alpha helical structure, and the antimicrobial/anticancer peptide generation instruction is: I would like to generate a protein that has antimicrobial/anticancer activity.

### 4.5 Protein Sequence Generation

During the generation process, prefix tokens encoded desired property were first fed into a prefix encoder to get task-specific prefix embeddings. Sequence generation proceeds autoregressively, where at each step, the subsequent amino acid was produced according to its preceding sequence context and the conditioned prefix embeddings.

To enhance the generative diversity and mitigate repetitive amino acids, a stochastic decoding strategy combining top-*k* sampling [37] with a repetition penalty mechanism [18] was employed. Specifically, at each step, the model’s output logits are truncated to retain only the top *k* most probable tokens. A temperature-scaled softmax operation then renormalized these probabilities, e.g. the sampling probability *p*_*i*_ of *i*-th token being sampled from the top *k* tokens was computed by Eq. (9):

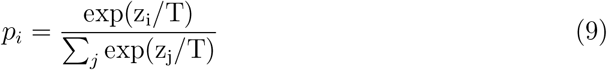

where *z*_*i*_ denotes the logit for *i*-th token, *T* is the temperature parameter controlling entropy. To further discourage repetitive outputs, a repetition penalty dynamically reduces the probabilities of tokens present in the previously generated sequence *g*. This is formalized by modifying the logits as Eq. (10):

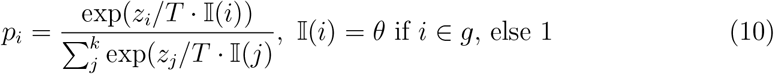

where *θ ∈* (0, 1) servers as a tunable penalty coefficient.

### 4.6 Evaluation Metrics

To comprehensively assessed the quality and validity of generated protein sequences, we employed a multi-dimensional evaluation framework, encompassing structural, energetic, functional, and sequence-level metrics.

**Structural validity** was valuated through three complementary approaches: (1) pLDDT (predicted Local Distance Difference Test), a confidence metric for predicted protein structure, was calculated using ESMFold, where higher average pLDDT scores (range: 0-100) suggest stable and plausible structures. (2) Secondary structure composition was quantified using the DSSP tool. Following the structural prediction via ESM-Fold [25], amino acid residues were labeled with the corresponding secondary structure property, including *α*-helix (H), Isolated *β*-bridge residue (B), Strand (E), 3-10 helix (G), *π*-helix (I), Turn (T), Bend (S) and other (-). In this study, we particularly focused on alpha-helix and beta-sheet content. (3) Ramachandran analysis was conducted to assess backbone dihedral angle (*ϕ, ψ*) distributions, highlighting energetically favorable and forbidden conformations. Rama-Z score quantifies deviations from preferred angle distributions. Typical values [*−*3, +3] indicate structural normality, and outlier beyond *±*3 suggest potential structural errors.

**Energetic stability** was evaluated using Rosetta Relax with the default Rosetta full-atom energy terms and weights in the PyRosetta [8]. To minimize the time needed for relax, the maximum number of iterations was set to 20. In generally, the total score (Rosetta Energy Units, REU) should typically fall within the range of −1 to −3 per residue [49], where lower Rosetta energy conformers correlate with more relaxed structures [41].

**Functional and physicochemical fidelity** were assessed via state-of-the-art predictors. The antimicrobial function of generated AMPs was predicted by leveraging the CAMP server [46]. The anticancer function of generated ACPs were predicted through AntiCP2.0 [1]. The physicochemical properties of generated AMPs and ACPs, including length, charge, charge density, hydrophobicity, hydrophobic moment, and isoelectric point, were calculated by using modlAMP [32]. Sequences with a length greater than 100 amino acids were removed before prediction.

**Sequence analysis** was also evaluated with four complementary approaches: (1) Sequence novelty measures how different the generated sequences are from the training data. It was measured through the follow formula [34].

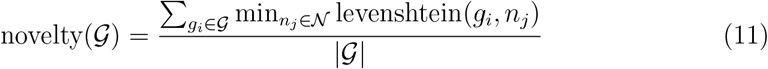

where *𝒢* denotes the set of generated protein sequences, and *𝒩* represents the set of natural proteins used in the training dataset. *g*_*i*_ is the *i*-th generated sequence, and *n*_*j*_ is the *j*-th natural sequence. A higher novelty score indicates that the designed proteins are more dissimilar from natural proteins, reflecting better generalization and less memorization. (2) Sequence diversity quantifies how different the generated sequences are from each other. It was calculated by averaging the Levenshtein edit distances across all pairwise combinations:

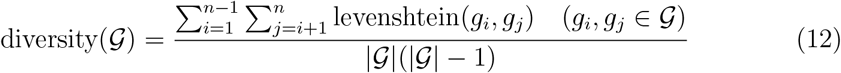

where *𝒢* denotes the set of generated protein sequences, *g*_*i*_, *g*_*j*_ are the *i*-th and *j*-th generated sequence. A higher diversity score suggests a broader exploration of the sequence space, which is desirable for generating varied and unique proteins. (3) Sequence-structure identity distribution measures the relationship between sequence and structure, where sequence identity was performed by BLAST, and the structure identity was measured by TM score via TM-align [51]. The low sequence identity and high TM-score compared to natural proteins indicates the generated proteins are novel and have functional activity. (4) Perplexity measures the uncertainty of sequence generation by calculating the exponential value of the cross-entropy loss. Lower perplexity values indicate better model performance, reflecting higher confidence in sequence generation. In this study, the perplexity were performed using the ProGen2-xlarge model.

### 4.7 Experimental Setup

To estimate the effectiveness of prefix tuning for controllable protein generation, we compared the propposed PrefixProt with several model variants: pre-trained Prot-GPT2, finetuned ProtGPT2, other PEFT-based ProtGPT2 (LoRA, (IA)^3^, and VeRA), and text-guided ProtLLMs (ProLLaMA and InstructProtein), as well as two baselines: the natural proteins in testing data and the randomly generated proteins. All models were trained for 100 epochs using AdamW optimizer [29]. The random generation was constructed by stochastically assembling amino acids according to their empirical background distribution in the training dataset.

For the alpha-helix structure dataset, the learning rates were set to 5e-6 for fine-tuned ProtGPT2, 5e-3 for PrefixProt, (IA)^3^, and VeRA, and 2e-4 for LoRA. The batch size for all models were set as 8. For antimicrobial and anticancer peptide datasets, the learning rates were set to 1e-6 for fine-tuned ProtGPT2, 1e-3 for PrefixProt, (IA)^3^, and VeRA, and 2e-4 for LoRA. The batch size for all models were set as 32. A polynomial learning rate decay schedule was applied across all tasks, decaying to a final rate of 1e-7 with a polynomial exponent of 3.

The prefix length was set to 100 virtual tokens for the alpha-helix proteins and 20 tokens for functional peptide generation. During inference, amino acids were sampled from top 500 predictions at each step, with repetition penalty of 1.2 to mitigate redundant sequences. To ensure a fair comparison, we generated 2,000 alpha-helix proteins, 1000 antimicrobial peptides, and 200 anticancer peptides by each compared method. In the prefix token combination experiments, all prefix tokens were retrained with length of 20 tokens. Two combination strategies were performed on the prefix matrices: element-wise averaging and direct concatenation.

For the pre-trained ProtGPT2 baseline, generation was constrained to a maximum length of 20 residues to align with the peptide-focused scope of our study, despite the model’s capacity to generate longer sequences. This restriction reflects the prevalence of short functional peptides in real-world applications and ensures parity in evaluation across methods.

## 5 Acknowledgements

This work is supported by the Natural Science Foundation of China (62102118), Guangdong Basic and Applied Basic Research Foundation (2025A1515010185), the Shenzhen Colleges and Universities Stable Support Program (GXWD20220811170504001), Shenzhen Science and Technology Program (JCYJ20230807094318038, KQTD2024072910215406), Futian Healthcare Research Project (FTWS2023091).

## Supplementary Files

### Figures

**Figure S1.**
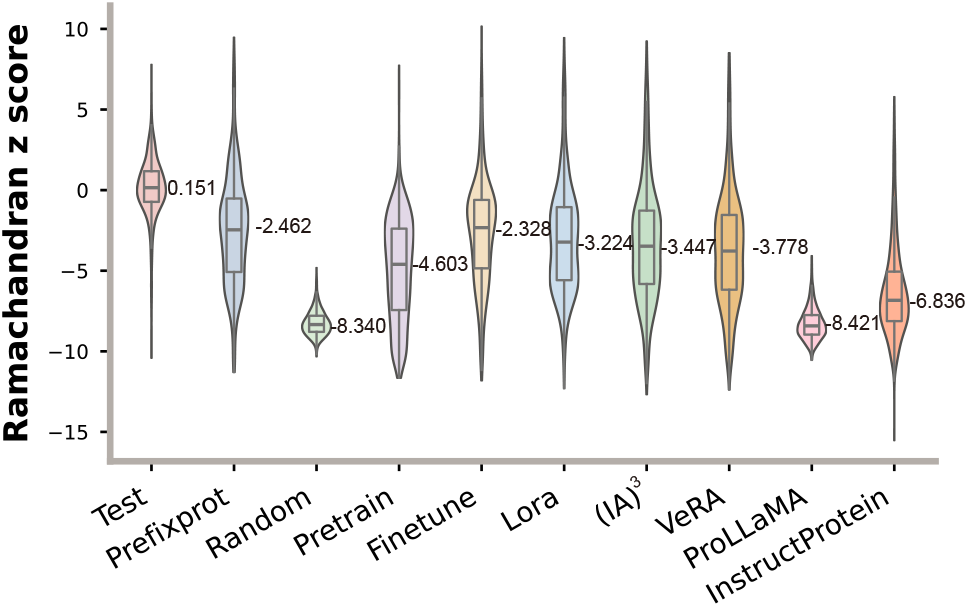
The distribution of Ramachandran z-score across different methods on alpha-helix protein generation.

**Figure S2.**
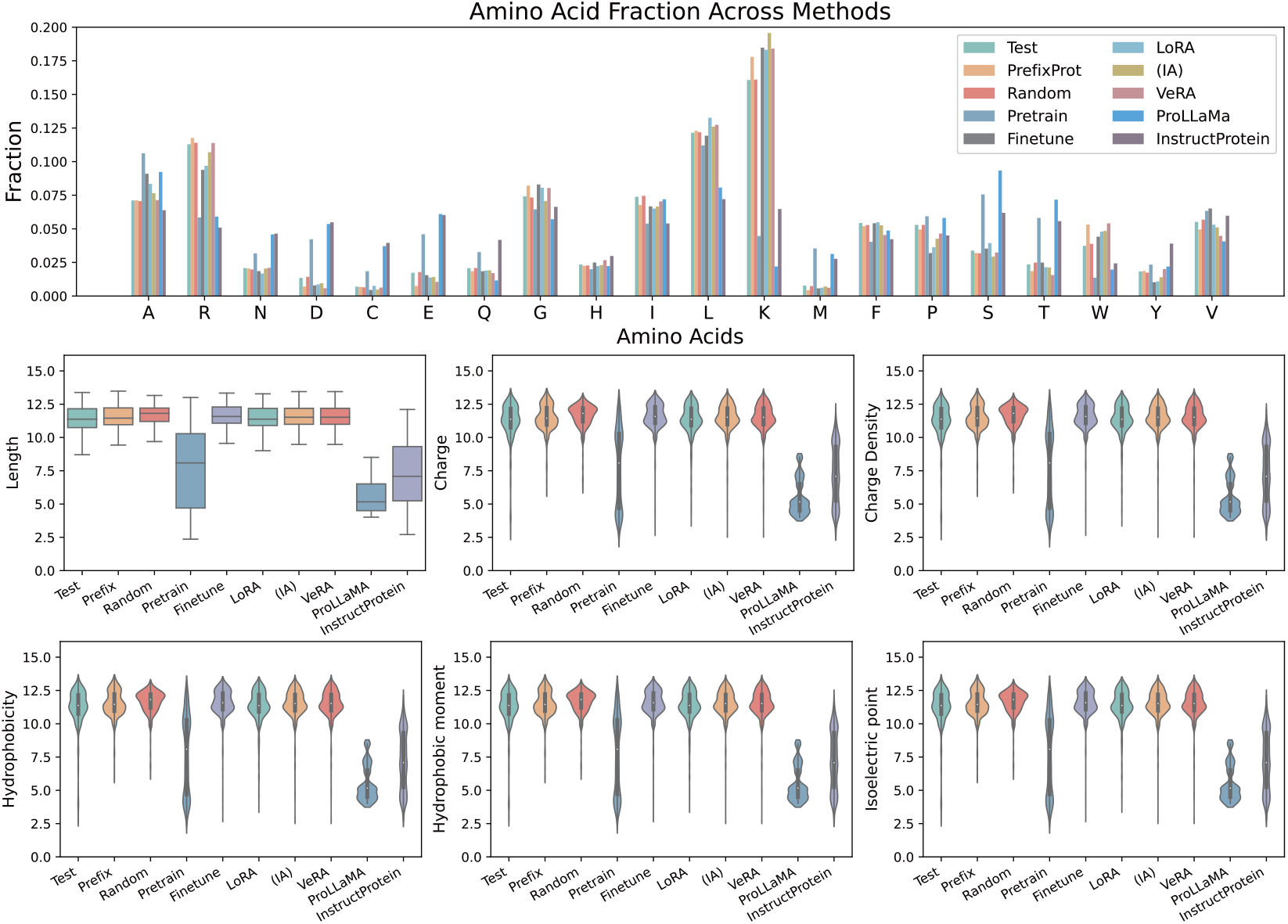
Physicochemical property comparison in generated AMPs.

**Figure S3.**
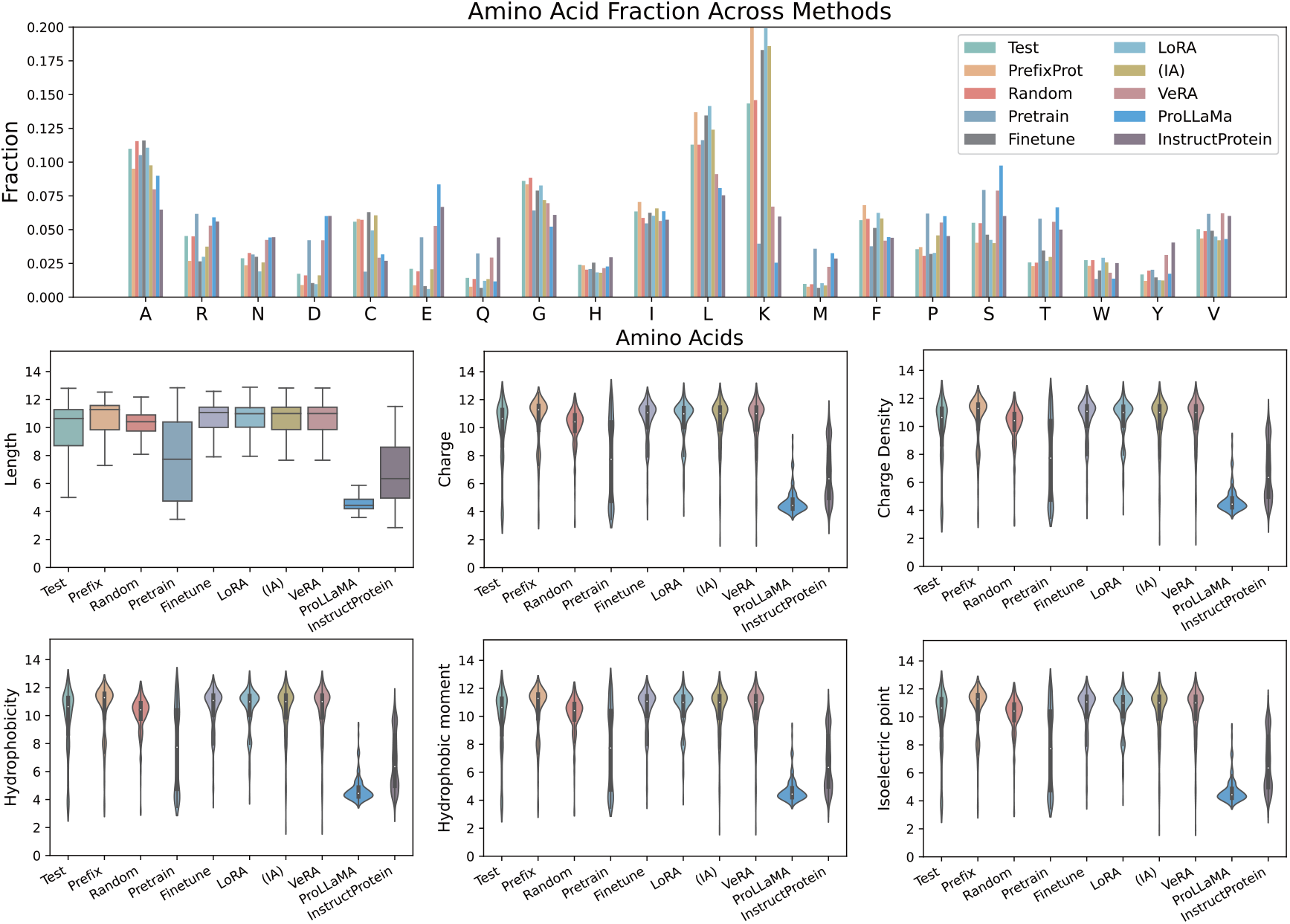
Physicochemical property comparison in generated ACPs.

**Figure S4.**
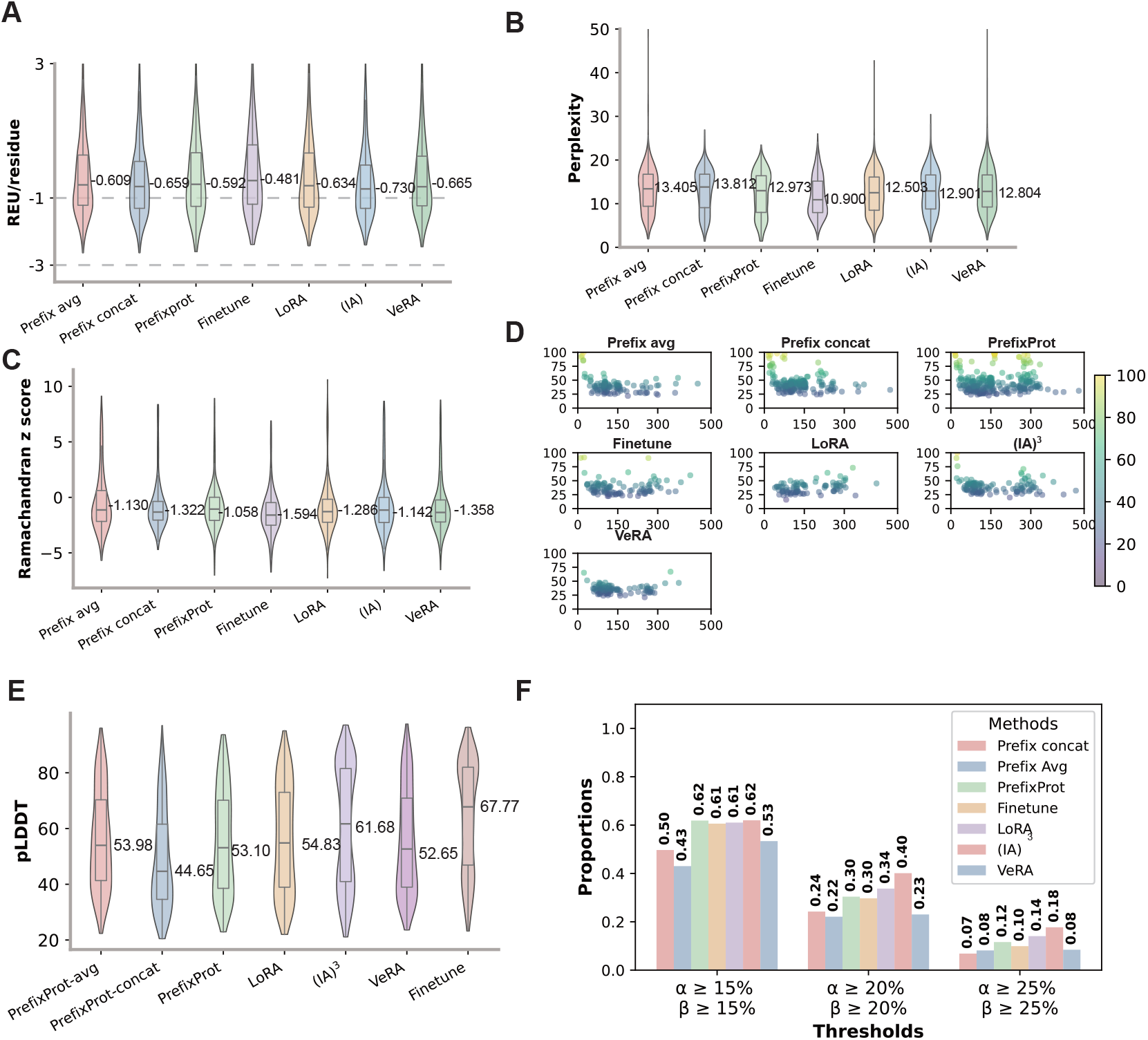
(A–C) The distribution of energy score (A), perplexity (B), and Ramachandran z-score (C) across different compared methods. (D) Pairwise sequence identity against a alpha-beta protein dataset. (E) pLDDT distribution of predicted protein structure trained on whole alpha proteins, beta proteins, and alpha-beta proteins. (F) Proportion of high-confidence generated proteins (pLDDT*>*70) with alpha-helix and beta-sheet content exceeding defined thresholds of 0.15, 0.20, and 0.25.

**Figure S5.**
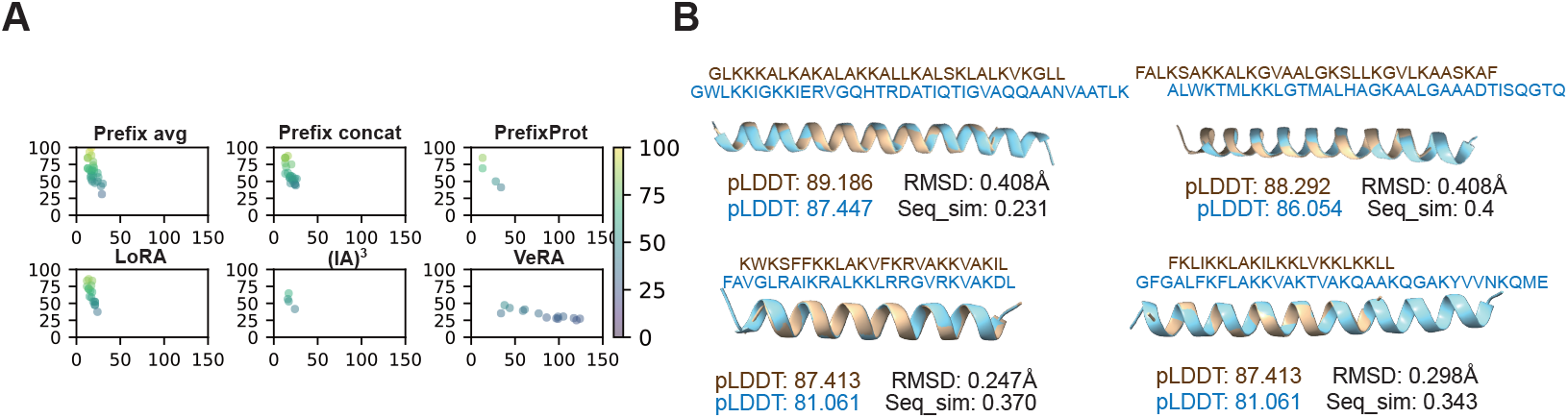
(A) Pairwise sequence identity against a multi-property protein dataset(AM-AC peptides). (B) The structure alignment of two samples generated using averaging (top) and concatenation (bottom) prefix combination approaches, where the brown and blue colors represent the natural and generated AM-AC peptides, respectively.

### Tables

**Table S1.**
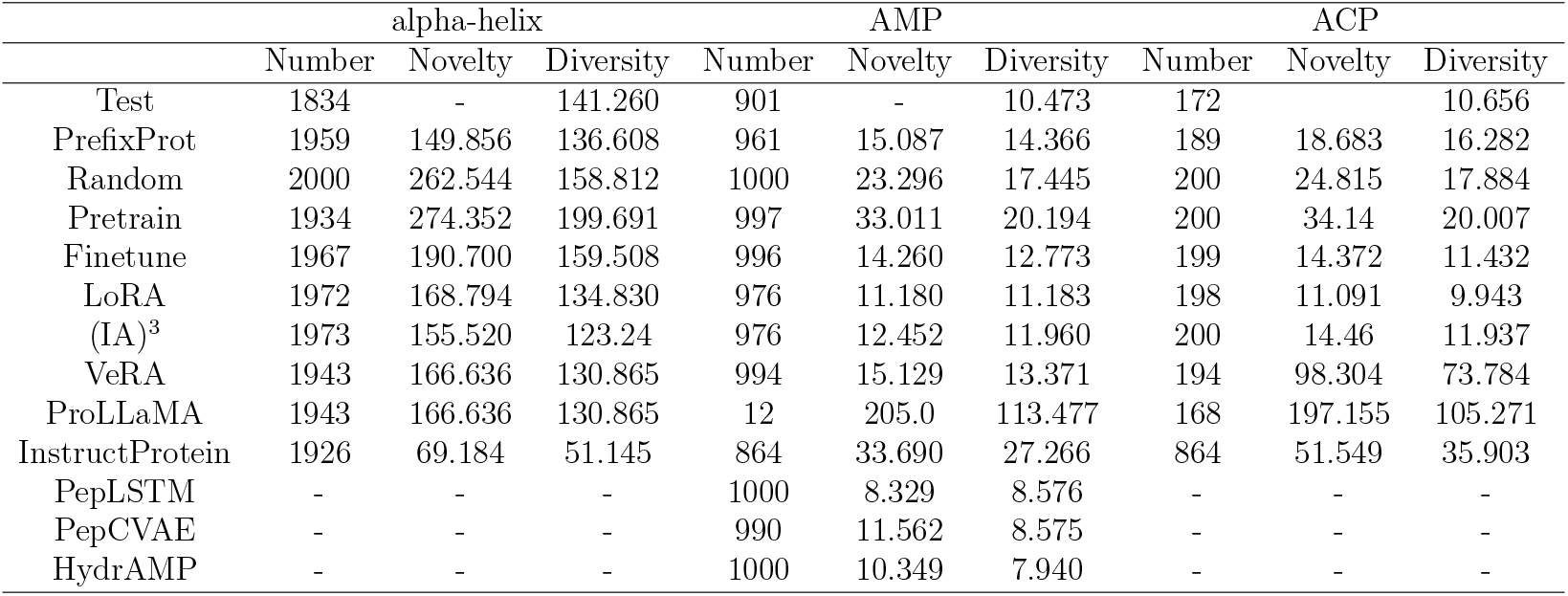
The performance of generated protein, including the number of proteins, mean perplexity, novelty and diversity.

|  | alpha-helix |  |  | AMP |  |  | ACP |  |  |
| --- | --- | --- | --- | --- | --- | --- | --- | --- | --- |
|  | Number | Novelty | Diversity | Number | Novelty | Diversity | Number | Novelty | Diversity |
| Test | 1834 | - | 141.260 | 901 | - | 10.473 | 172 | - | 10.656 |
| PrefixProt | 1959 | 149.856 | 136.608 | 961 | 15.087 | 14.366 | 189 | 18.683 | 16.282 |
| Random | 2000 | 262.544 | 158.812 | 1000 | 23.296 | 17.445 | 200 | 24.815 | 17.884 |
| Pretrain | 1934 | 274.352 | 199.691 | 997 | 33.011 | 20.194 | 200 | 34.14 | 20.007 |
| Finetune | 1967 | 190.700 | 159.508 | 996 | 14.260 | 12.773 | 199 | 14.372 | 11.432 |
| LoRA | 1972 | 168.794 | 134.830 | 976 | 11.180 | 11.183 | 198 | 11.091 | 9.943 |
| (IA)<sup>3</sup> | 1973 | 155.520 | 123.24 | 976 | 12.452 | 11.960 | 200 | 14.46 | 11.937 |
| VeRA | 1943 | 166.636 | 130.865 | 994 | 15.129 | 13.371 | 194 | 98.304 | 73.784 |
| ProLLaMA | 1943 | 166.636 | 130.865 | 12 | 205.0 | 113.477 | 168 | 197.155 | 105.271 |
| InstructProtein | 1926 | 69.184 | 51.145 | 864 | 33.690 | 27.266 | 864 | 51.549 | 35.903 |
| PepLSTM | - | - | - | 1000 | 8.329 | 8.576 | - | - | - |
| PepCVAE | - | - | - | 990 | 11.562 | 8.575 | - | - | - |
| HydrAMP | - | - | - | 1000 | 10.349 | 7.940 | - | - | - |

**Table S2.** The settings of STREME for finding the motif of the generated candidate multi-functional peptide, and the peptide with both antibacterial and anticancer functions.

| Parameter | value |
| --- | --- |
| Minimum width | 3 |
| Maximum width | 8 |
| $p$ value threshold | 0.05 |
| Number of motifs | 10 |
| Align sequences on their | Centers |

